# Stable coexistence of *Citrobacter rodentium* with a lytic bacteriophage during *in vivo* murine infection

**DOI:** 10.1101/2025.07.03.662908

**Authors:** Audrey Peters, Hiba Shareefdeen, Julia Sanchez-Garrido, Eli J. Cohen, Rémi Denise, Joshua L C Wong, Morgan Beeby, Colin Hill, Gad Frankel

## Abstract

Bacteriophages are ubiquitously present in bacterial communities, yet phage-bacteria interactions in complex environments like the gut remain poorly understood. While antibiotic resistance is driving a renewed interest in phage therapy, most studies have been conducted in *in vitro* systems, offering limited insight into the complexity of such dynamics in physiological contexts. Here, we use *Citrobacter rodentium* (CR), a natural mouse-restricted enteric pathogen and well-established model for human enteropathogenic and enterohaemorrhagic *Escherichia coli* (EPEC and EHEC) infections, to investigate phage-pathogen interactions *in vivo*. We isolate and characterise Eifel2, a novel lytic phage infecting CR, and generate anti-phage specific antibodies that enable the visualisation of phage infections *in vitro*. In a murine model of CR infection, oral administration of Eifel2 led to robust phage replication in the gut without reducing the bacterial burden or infection-associated inflammation, confirming the establishment of a stable coexistence in the gut. Despite the emergence of a sub-population of phage-resistant CR mutants *in vivo*, they did not undergo clonal expansion, indicating that additional selective pressures impaired their widespread dissemination in the gut. Together, our findings demonstrate that imaging approaches can capture key infection stages *in vitro*, while *in vivo* models are essential for capturing the complexity of phage-bacteria interactions. This work highlights the importance of studying phage therapy in host-pathogen contexts that include a normal microbiota and a suitable host environment, where dynamic co-existence rather than eradication may define therapeutic outcomes.

**Importance:** Bacteriophages, or phages, are viruses which can either kill or persist inside bacteria. Current interests in phage biology are in part ignited by the fact that they could be used to treat infections caused by antibiotic-resistant bacteria. However, most of our understanding of phage-bacterial interactions comes from in vitro models and/or in vivo gut models relying on altering the endogenous microbiota. Here we report the finding of a novel phage, Eifel2, which specifically targets *Citrobacter rodentium* (CR), the mouse equivalent of human diarrhoeagenic *E. coli* pathogens. Despite effectively killing CR in vitro, CR and Eifel2 develop a coexistence relationship in mice with an intact microbiota. While CR phage-resistant mutants emerge, host and microbial factors constrain their expansion. This work highlights the importance of studying phage therapy in host-pathogen contexts that include the complete microbiota, where therapeutic outcomes may rely on dynamic co-existence and containment rather than eradication.

## Introduction

Lytic bacteriophages or phages, viruses that specifically infect and lyse bacteria, are receiving renewed interest as an alternative to antibiotics, particularly to treat multidrug-resistant bacterial pathogens ^1–3^. However, translating promising *in vitro* results into successful therapeutic outcomes *in vivo* remains a challenge ^2,4^. Recent studies in murine models have shown that introducing phages can lead to stable coexistence between phages and their bacterial hosts in the gut instead of bacterial eradication ^5,6^. Such systems support a dynamic equilibrium, with stable levels of both phage and bacteria over time. This suggests active phage replication, potentially driven by protected bacterial replication niches in the gut ^7^, phage host-range expansion to microbiota members ^8^, or bacterial and phage coevolution leading to resistant subpopulations ^9,10^. Resistance can arise from mutation of phage receptors and often involves modification of surface structures such as lipopolysaccharide (LPS) or outer membrane proteins, which can carry fitness trade-offs *in vivo*, and thus do not lead to a predominantly phage-resistant population ^6^.

One of the major limitations to study phage-bacteria interactions in the gut is the availability of physiologically relevant infection models. Mice with an intact microbiota are usually refractory to exogenous bacterial infection, and therefore antibiotic pre-treatment or other methods to interfere with colonisation resistance are needed ^11^. This limits our ability to investigate phage-bacteria interactions and key processes such as the emergence and fitness cost of phage resistance in translational models, where they can be profoundly influenced by factors such as physicochemical conditions, host immune responses, spatial structure and microbial community dynamics.

To overcome these limitations, *Citrobacter rodentium* (CR), a natural murine enteric pathogen, has been widely used as a surrogate model for EPEC and EHEC infections *in vivo*. CR, shares key infection mechanisms with EPEC and EHEC, including the formation of attaching and effacing (A/E) lesions. CR, which robustly colonises mice with an intact microbiota, triggers colonic crypt hyperplasia (CCH), and gut inflammation, manifested by increased faecal levels of lipocalin-2 (LCN-2) ^12–14^. As such, the CR model provides a valuable opportunity to study, not only phage-bacteria interactions *in vivo*, but also the broader gut ecological and immunological dynamics that shape phage therapy outcomes in a physiologically representative context. Despite this potential, relatively few phages infecting CR have been isolated to date ^15–18^, and their application in murine models of enteric infection remain unexplored.

In this study, we aimed to isolate and characterise novel lytic phages that infect CR. We investigated phage-bacteria interactions *in vitro* and *in vivo* by combining immunofluorescence imaging with our well-established mouse model of CR infection.

## Results

### Characterisation of CR lytic phages Eifel1 and Eifel2

Phage Eifel1 infecting CR strain ICC169 was isolated from sewage water samples collected in the Eifel region in Germany. Eifel1 forms small, translucent plaques (<1 mm in diameter) on soft agar CR lawns (Fig. 1A). During the purification process, we also recovered larger plaques (1.5–2 mm), appearing among the Eifel1 plaques (Fig. 1B). We isolated these separately and designated the new phage variant Eifel2.

**Figure 1.**
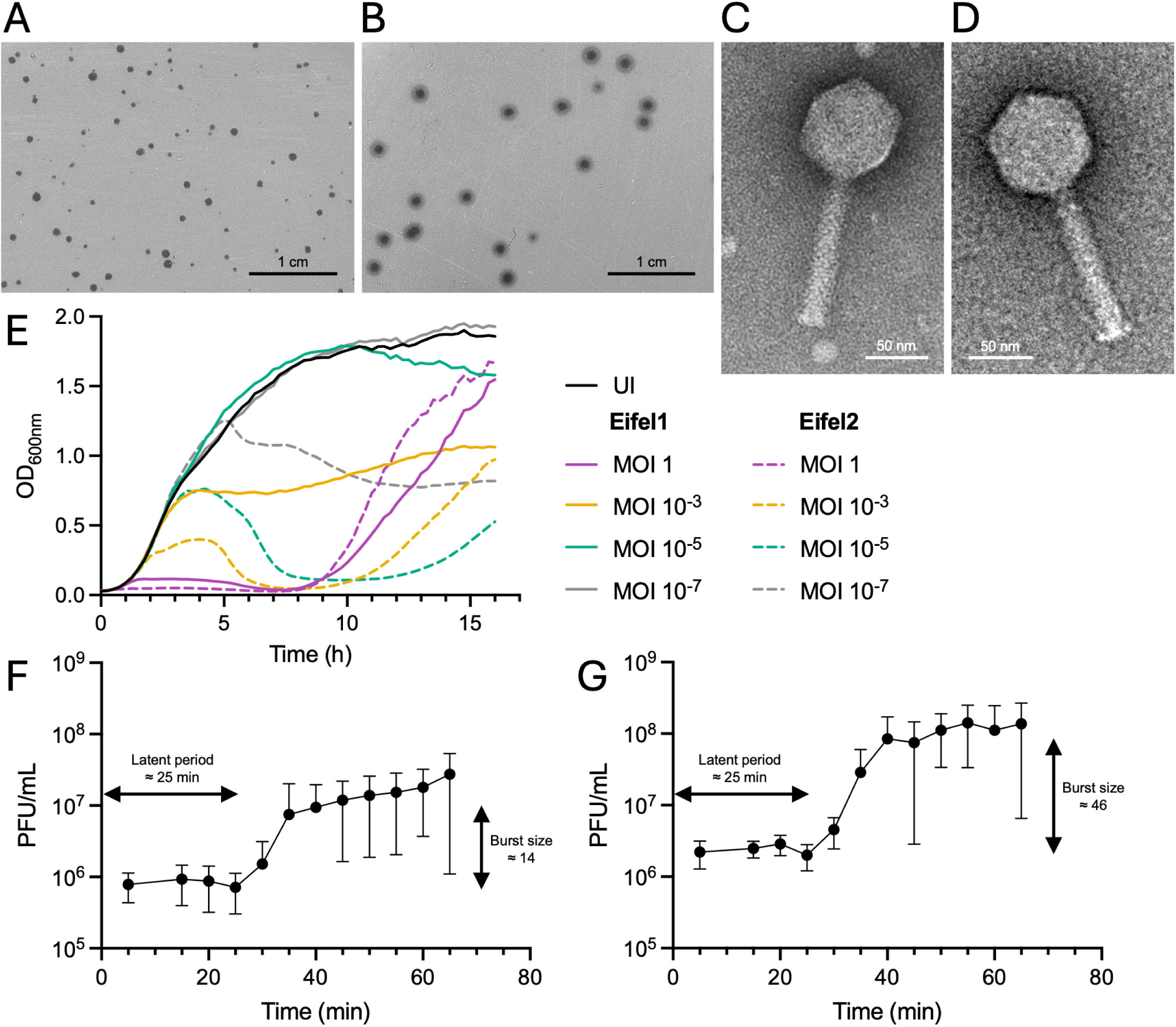
Morphology and replication characteristics of phages Eifel1 and Eifel2. (A-B) Plaque morphology of phages Eifel1 (A) and Eifel2 (B) on a double layer agar (DLA) plate with CR. Scale bar: 1 cm. (C-D) Representative transmission electron micrographs of uranyl acetate negatively stained Eifel1 (C) and Eifel2 (D) virions. Scale bar: 50 nm. (E) Lysis kinetics of CR cultures infected with Eifel1 (solid lines) and Eifel2 (dashed lines) at the indicated MOIs or uninfected (UI) (solid black line). Data are shown as mean of n = 3 independent biological repeats. (F-G) One-step growth curve of phages Eifel1 (F) and Eifel2 (G) in CR at a MOI of 1, showing latent period and burst size. Phages were allowed to adsorb to CR for 5 min, after which unbound phages were washed off, and phage titres were determined at indicated timepoints. Data are shown as mean ΢ standard deviation (SD) of n = 5 independent biological repeats.

Transmission electron microscopy (TEM) revealed that Eifel1 and Eifel2 share a similar morphology, with virions formed of an icosahedral head and a long contractile tail (Fig. 1C, D). The head of Eifel1 measured 76.2 ± 2.8 nm (mean ± SD) in length and 73.8 ± 2.9 nm in width, while the tail measured 110.9 ± 4.5 nm and the neck 11.0 ± 2.5 nm (n = 23 virions). Eifel2 exhibited comparable dimensions: head length of 77.8 ± 3.0 nm, width of 74.2 ± 3.4 nm, tail length of 116.2 ± 9.2 nm, and neck size of 11.5 ± 2.5 nm (n = 56 virions). Head width and neck length showed no significative differences between Eifel1 and Eifel2 (*P* > 0.05 by Mann-Whitney test), whereas head length and tail length were significantly longer in Eifel2 (*P* = 0.0176 and *P* = 0.0005 by Mann-Whitney test respectively).

To assess their lytic activity, we infected CR with Eifel1 and Eifel2 at multiplicities of infection (MOIs) ranging from 1 to 10⁻⁷ and monitored optical density (OD600) for 16 h (Fig. 1E and Fig. S1A, B). For most MOIs tested, Eifel1 and Eifel2 reduced OD_600_ following an initial growth period of 2–10 h, with shorter delays at higher MOIs. After this initial decrease, indicating bacterial lysis, OD_600_ levels increased again, suggesting the emergence of resistant bacteria. Notably, cultures infected with Eifel2 at MOI 1 showed no growth until 8 h post-infection (Fig 1E). At the lowest MOIs (10⁻⁶ and 10⁻⁷), Eifel1 had no observable impact on bacterial growth compared to uninfected controls. Across all MOIs tested, Eifel2 caused a more rapid and pronounced OD_600_ decrease than Eifel1, suggesting higher lytic activity.

To investigate whether the difference in infection kinetics was caused by variations in burst size or infection cycle length, we conducted one-step growth curve analyses. Both Eifel1 and Eifel2 exhibited a latent period of 25 min. However, Eifel1 released an average of 14 new virions per infected cell (Fig. 1F), while Eifel2 released 46 virions per infected cell (Fig. 1G), indicating a significantly higher burst size (*P* < 0.05 determined by lognormal Welch’s *t* test). During the latent period, Eifel1 maintained a concentration of ∼8.25 × 10⁵ PFU/mL, whereas Eifel2 reached ∼2.39 × 10⁶ PFU/mL, suggesting a higher adsorption rate for Eifel2.

We also tested the host range of Eifel1 and Eifel2 against 21 Gram-negative strains (3 *Citrobacter* species, 11 *Escherichia coli* strains, 2 *Klebsiella* species, *Enterobacter cloacae, Pseudomonas aeruginosa*, *Salmonella enterica* and *Shigella sonnei*); we also attempted to infect a Gram-positive *Staphylococcus aureus* strain. None of the tested isolates were susceptible to either Eifel1 or Eifel2 infection, highlighting the specificity of both phages for CR. Thus, we concluded that Eifel1 and Eifel2, whilst having been isolated from the same origin and sharing physical morphology and host range, showed a significant difference in their lytic efficiency.

### Comparative genomic analysis of Eifel1 and Eifel2

To better understand the relationship between Eifel1 and Eifel2, their genomes were sequenced following phage purification. Assembly of Eifel1 revealed that its genome consists of a linear double-stranded DNA of 88,123 base pairs (bp) with a GC content of 39.1%. We identified 147 open reading frames (ORFs), of which 57 were assigned putative functions, along with 21 tRNA genes (Fig. 2A). We found no genes encoding integrases or other lysogeny-associated elements, and PhageAI predicted a lytic lifestyle with 99.97% confidence.

**Figure 2.**
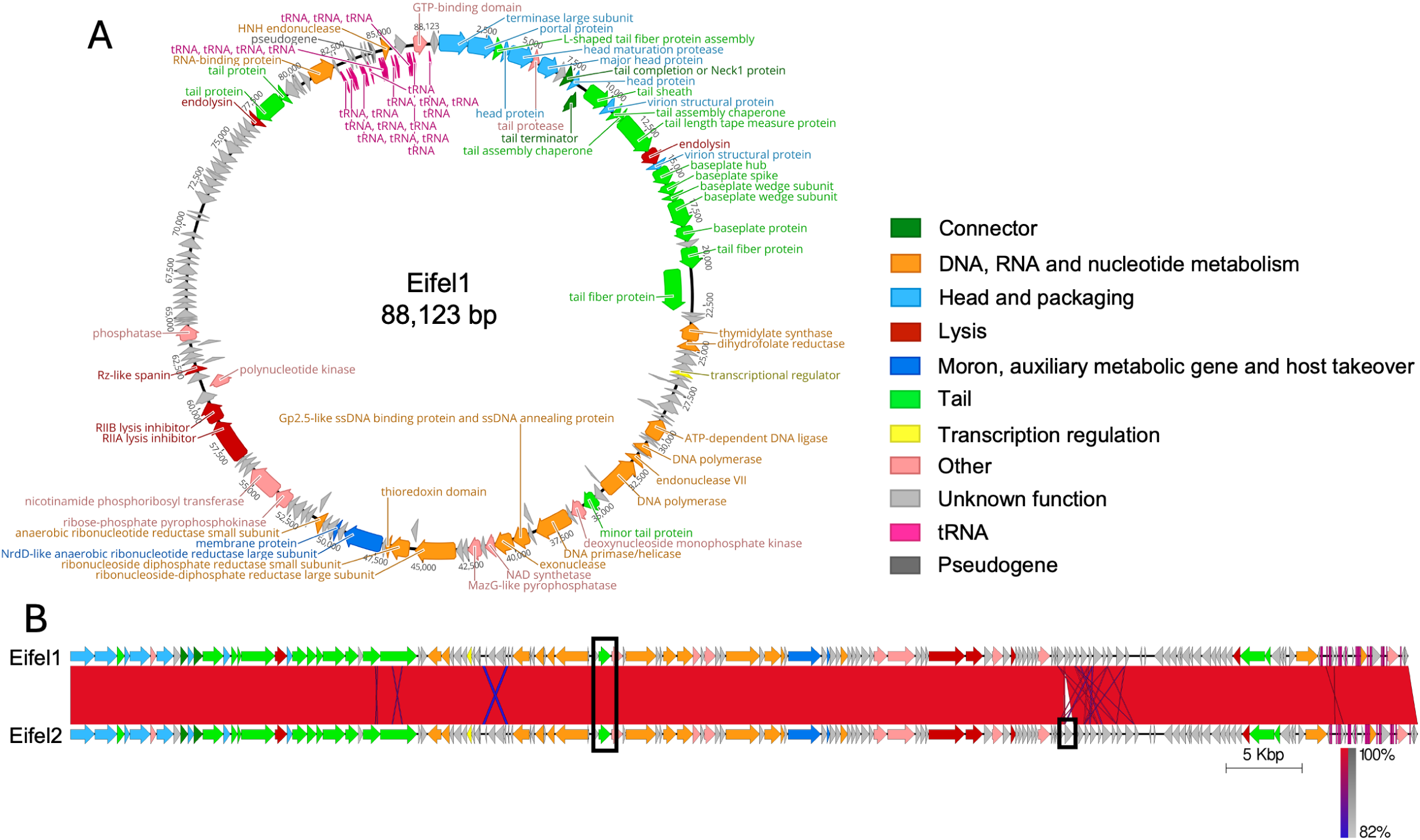
Genomic organisation and comparison of phages Eifel1 and Eifel2. (A) Circular map of the Eifel1 genome, annotated based on ORFs predicted using Phold (https://github.com/gbouras13/phold) and displayed as directional arrows. ORFs are coloured according to putative function, with predicted gene products indicated. (B) Whole genome alignment of Eifel1 and Eifel2 performed using the progressive Mauve algorithm ^19^ and visualised with Easyfig^20^. ORFs are represented as arrows and coloured as in panel A. Genomic differences between the two phages are highlighted by black boxes.

Taxonomic classification using taxMyPhage placed Eifel1 as a novel species in the *Felixounavirus* genus (subfamily *Ounavirinae*, family *Andersonviridae*, class *Caudoviricetes*) in line with the most recent taxonomy update from the International Committee on Taxonomy of Viruses (ICTV). According to the latest ICTV Master Species List (MSL40.v1), the *Felixounavirus* genus currently comprises 97 recognised species. To further support this classification, we constructed a phylogenetic tree using VICTOR, incorporating the representative genome of each *Felixounavirus* species alongside the Eifel1 genome. The resulting tree confirmed that Eifel1 forms a distinct clade, supporting its classification as a novel species within the genus (Fig. S2A).

Eifel2 has a linear double-stranded DNA genome of 88,736 bp and PhageAI predicted a lytic lifestyle at 99.96% confidence. Whole-genome alignment of Eifel1 and Eifel2 revealed almost identical sequences, supporting the classification of Eifel2 within the *Felixounavirus* genus as a member of the same novel species as Eifel1 (Fig. S2A). Only two differences were identified between the two genomes, and these were confirmed by Sanger sequencing (Fig. 2B). The first was a 12 bp deletion in an Eifel2 gene encoding a minor tail protein, which results in the loss of four amino acids (aa) in the predicted protein (Fig. S2B). Structural predictions generated with AlphaFold3 suggested that this deletion would cause a loop in an α-helix at the C-terminal domain of the protein (Fig. S2C). Interestingly, BLAST analysis of the Eifel2 minor tail protein revealed homologous sequences with none containing the 4 aa deletion. The second difference was the presence of an additional 513 bp ORF in Eifel2, encoding a putative protein of unknown function (Fig. S2D). This gene showed no similarities with the CR encoded genes or endogenous plasmids. However, homologous sequences were detected in other *Felixounavirus* phage genomes within GenBank, such as *Salmonella* phage FelixO1 (accession number: AF320576) (Fig. S2E).

### Tracking Eifel2 infection dynamics by immunofluorescence microscopy

Considering that Eifel1 and Eifel2 differ by only one 4 aa deletion and the acquisition of one 513 bp ORF, we selected the latter for further characterisation.

To investigate the interaction between Eifel2 and CR, we generated anti-sera to Eifel2. We immunised mice with purified Eifel2 phage particles according to the schedule in Fig. 3A, with control mice receiving a no phage buffer control. Dot blot assays were performed using Eifel2, Eifel1, and two controls: (i) filtered supernatant from a CR overnight culture, and (ii) an additional tailed phage isolated during this study infecting CR named ColRes, and belonging to a novel species within the *Tequatrovirus* genus (plaque morphology and TEM image are shown in Fig. S3A and S3B, respectively). Serum from immunised mice detected both Eifel1 and Eifel2, whereas no signal was observed using CR supernatant and only a faint signal was detected for ColRes. No reactivity was seen with serum from control animals, indicating the generated antibodies specifically targeted Eifel2, and cross-reacted with Eifel1 (Fig 3B).

**Figure 3.**
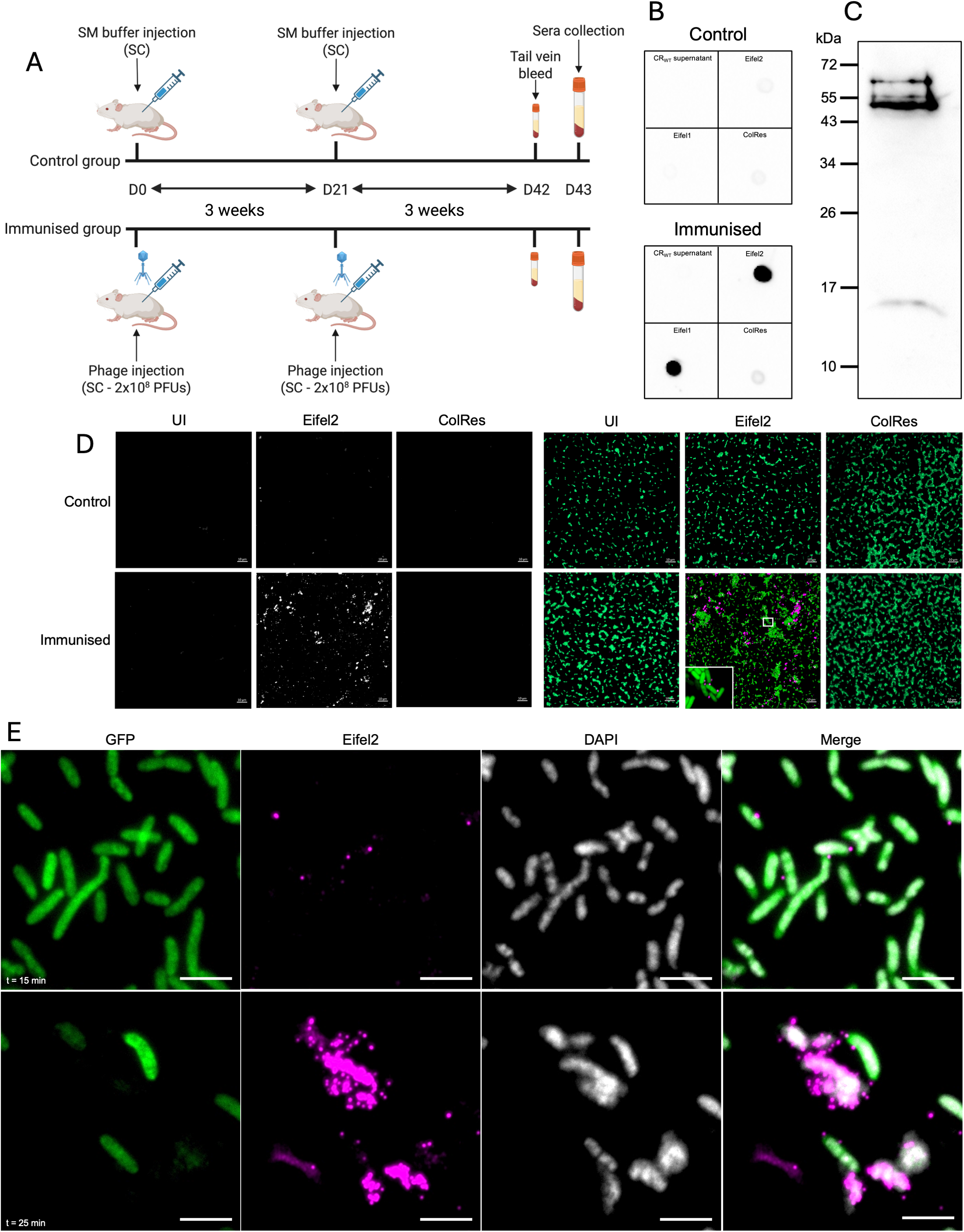
Generation and application of anti-Eifel2 antibodies in immunofluorescence microscopy. (A) Schematic representation of the mouse immunisation protocol used to generate serum reactive to Eifel2T. (B) Dot blot showing reactivity of serum from mock-immunised mice (top panel) or immunised mice (bottom panel) to CR overnight culture filtered supernatant and phages Eifel2, Eifel1 and ColRes. (C) Western blot analysis of purified Eifel2 lysed virions probed with anti-Eifel2 serum from immunised mice. (D) Immunofluorescence detection of phage Eifel2 particles. GFP-expressing CR were either uninfected (left column), infected with Eifel2 (middle column), or infected with ColRes (right column) for 5 min. Samples were stained using either mock-immunised serum (top row) or anti-Eifel2 immunised serum (bottom row). Left panel shows phage-specific channel (white); right panel shows merged images of CR signal (GFP, green) and phage signal (magenta). A selected region is highlighted with a white box and shown at higher magnification to visualise phage binding at the single-cell level. Scale bar: 10 μm. (E) Time-course immunofluorescence analysis of Eifel2 infecting CR. GFP-expressing CR cultures were infected with Eifel2, fixed and stained at 15 min (top row) and 25 min (bottom row) post-infection. Columns show individual fluorescence channels: CR (GFP, green), phage (magenta), DNA (DAPI, white), and a merged image of all channels. Phage detection was performed using anti-Eifel2 serum. Scale bar: 5 μm. (B, C, D) Images are representative of n = 3 independent biological repeats. (E) Images are representative of n = 2 independent biological repeats.

ELISA revealed a ∼1000-fold increase in Eifel2-specific IgG titres in the immunised group compared to controls (Fig. S3C). Western blot analysis of lysed Eifel2 virions revealed five distinct immunoreactive bands (Fig. 3C), indicating that the generated antibodies recognise multiple phage proteins.

We then used the anti-Eifel2 serum to follow phage binding and release by immunofluorescence microscopy (IF). To this end, we first infected CR expressing green fluorescent protein (GFP) with Eifel2. Cultures were fixed 5 min post-infection and stained with the serum of immunised or control mice. Uninfected bacteria and cultures incubated with ColRes were used as negative controls. No signal was detected for any of the samples stained with serum from mock-immunised mice. For samples stained with immunised serum, IF microscopy revealed strong fluorescent signals associated with Eifel2-infected bacteria (Fig. 3D), while no signal was detected in either control.

We next performed time-course infections using a one-step growth curve setup. Infected cultures were sampled at defined timepoints, fixed, and stained with serum from immunised mice for IF analysis (Fig. 3E). At 15 min post-infection, discrete dot signals are visible around CR, indicative of phage attachment. After 25 min, in line with the one-step growth curve data, we observed intense phage staining surrounding bacteria that displayed a reduced GFP signal and intense DAPI signal, consistent with bacterial lysis and phage release.

### Eifel2 coexists with CR in a murine model

We next aimed to study CR – Eifel2 interactions in vivo. To this end, we first assessed its stability across different temperatures and pH values. Phage titres remained stable at 4°C and 37°C for at least 24 h (Fig. S4A). In contrast, higher temperatures progressively reduced phage viability, and no plaques were detectable after 5 min incubation at 90°C. Freeze thawing phage preparations also led to a gradual decline in titre over time. Eifel2 titres were undetectable after 24 h of incubation at 37°C in buffers at pH 1, 2, 3 and 13, but titres remained stable for at least 24 h in buffers ranging from pH 4 to 12, indicating the phage can withstand the acidic pH in the gastrointestinal tract of fasted mice ^21^(Fig. S4B).

We next investigated phage-bacteria interactions in vivo by oral administration of Eifel2 to C57BL/6 mice. Mice infected with CR received daily phage treatment (1 x 10^9^ PFUs) by oral gavage, or SM buffer as a control, from 3 to 7 days post-infection (dpi), corresponding to the expansion phase, which is characterised by the spread to the colonic mucosa from the caecal patch and rapid bacterial proliferation ^22^. A non-infected group received phage treatment alone (Fig. 4A). We followed temporal bacterial and phage faecal shedding, until no phage was detectable. No differences were observed in CR shedding between phage-treated and untreated mice, with both groups shedding between 10⁸ and 10⁹ CFU/g of stool until 13 dpi, after which bacterial loads gradually declined to 10³–10⁴ CFU/g by 23 dpi (Fig. 4B).

**Figure 4.**
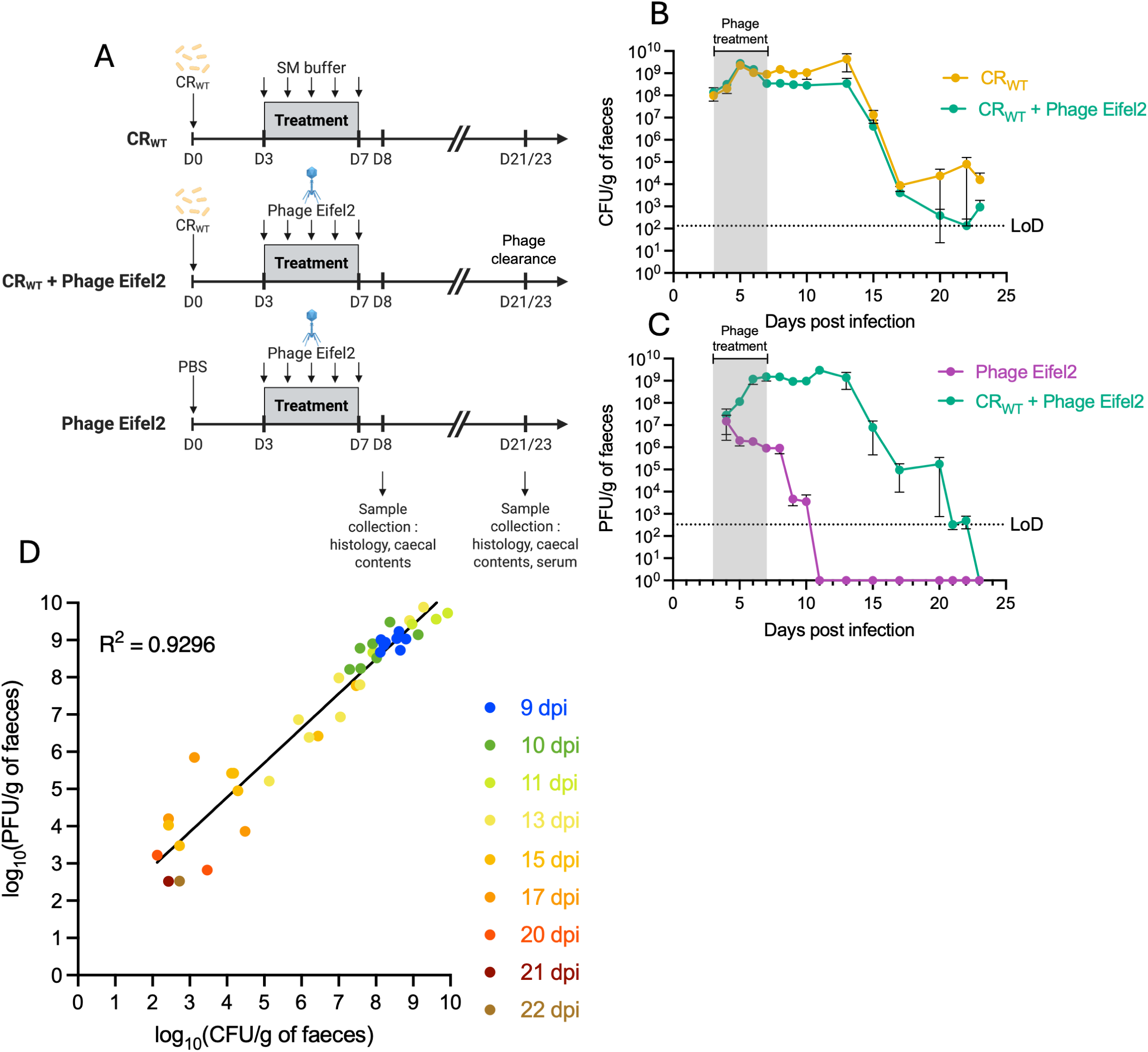
Eifel2 replicates in a murine model of CR infection without reducing bacterial burden. (A) Schematic representation of the mouse phage treatment experiment. Mice were orally infected with CR or mock-infected and received daily oral gavage of either Eifel2 phage suspension or SM buffer (mock treatment) from 3 to 7 dpi. Bacterial and phage shedding in faeces was monitored until phage clearance, which occurred at 21 and 23 dpi in two independent experiments. Mice were culled at 8 dpi and at the point of phage clearance. (B) Temporal quantification of CR shedding in faeces, expressed as CFU per g of faeces, in infected mice receiving phage treatment or mock treatment. From 0 to 8 dpi, sample sizes were n = 19 (untreated) and n = 20 (phage-treated) across 3 independent experiments. After 8 dpi, data are from n = 7 (untreated) and n = 8 (phage-treated) across 2 independent experiments. (C) Temporal quantification of Eifel2 shedding in faeces, expressed as PFU per g of faeces, in infected or mock-infected mice receiving phage treatment. For UI mice, sample sizes were n = 16 (0-8 dpi) and n = 8 (after 8 dpi) across 2 independent experiments. For infected mice, samples sizes were n = 20 (0-8 dpi) and n = 8 (after 8 dpi) across 3 independent experiments. (D) Correlation analysis of log_10_-transformed PFU and CFU counts in faeces of phage-treated infected mice. Each point represents values from an individual mouse at a specific timepoint after end of phage treatment (9-22 dpi). Simple linear regression is shown with the corresponding equation and R^2^. Data points below the LoD for either PFU or CFU were excluded. (B, C) Data are shown as mean standard error of the mean (SEM), and values below the limit of detection (LoD) were set to 1 for plotting.

Phage shedding, however, differed between CR-infected and uninfected animals. In both groups, faecal phage titres were 10⁷–10⁸ PFU/g one day after beginning of phage treatment. In uninfected mice, phage titres declined steadily to 10^6^ PFU/g despite continued treatment and became undetectable by 11 dpi (4 days after the end of the treatment). In contrast, CR-infected mice showed increasing phage titres that peaked at 10⁹ PFU/g of stool and remained elevated until 13 dpi (6 days after the end of the treatment). After this point, phage titres declined in parallel with CR loads and were undetectable by 23 dpi (Fig. 4C). CR and phage titres in the caecum at 8 dpi were consistent with faecal levels, with CR counts averaging ∼10^9^ CFU/g of caecal contents for both infected groups, although the phage-treated group exhibited slightly lower bacterial counts. Phage titres were ∼10⁸ PFU/g in infected mice and ∼10⁶ PFU/g in the phage-only group (Fig. S4C, D).

The observed differences suggest that phage replication occurs only in the presence of CR, and that oral administration of Eifel2 alone does not result in sustained colonisation due to the absence of a suitable commensal host. Correlation analysis of PFU and CFU counts for each mouse across different timepoints revealed a strong positive relationship between phage and bacterial levels in the gut (R² = 0.9296) (Fig. 4D), indicating a highly dynamic interaction *in vivo*. These findings confirm that Eifel2 can infect CR *in vivo*, thus maintaining high titres over time, and support the establishment of a stable co-existence between Eifel2 and CR within the gut environment.

### Phage – CR coexistence does not affect host – pathogen interactions

To visualise phage – CR interactions in the gut, distal colon sections were stained using the Eifel2 antiserum alongside anti-CR serum at 8 dpi. While no phage signal was detected, comparable levels of attached CR were observed in both phage-treated and untreated groups (Fig. 5A). Moreover, evaluation of colon histology samples revealed that both infected groups, regardless of phage treatment, exhibited CCH (crypt length ∼200 μm) at 8 dpi and at the time of phage clearance (21-23 dpi), while uninfected controls showed normal crypt lengths (∼150 μm) (Fig. 5B). As CCH is a hallmark of CR infection that reflects activation of the damage repair response and subsequent cell proliferation and crypt elongation, phage treatment does not appear to impact CR-induced barrier damage.

**Figure 5.**
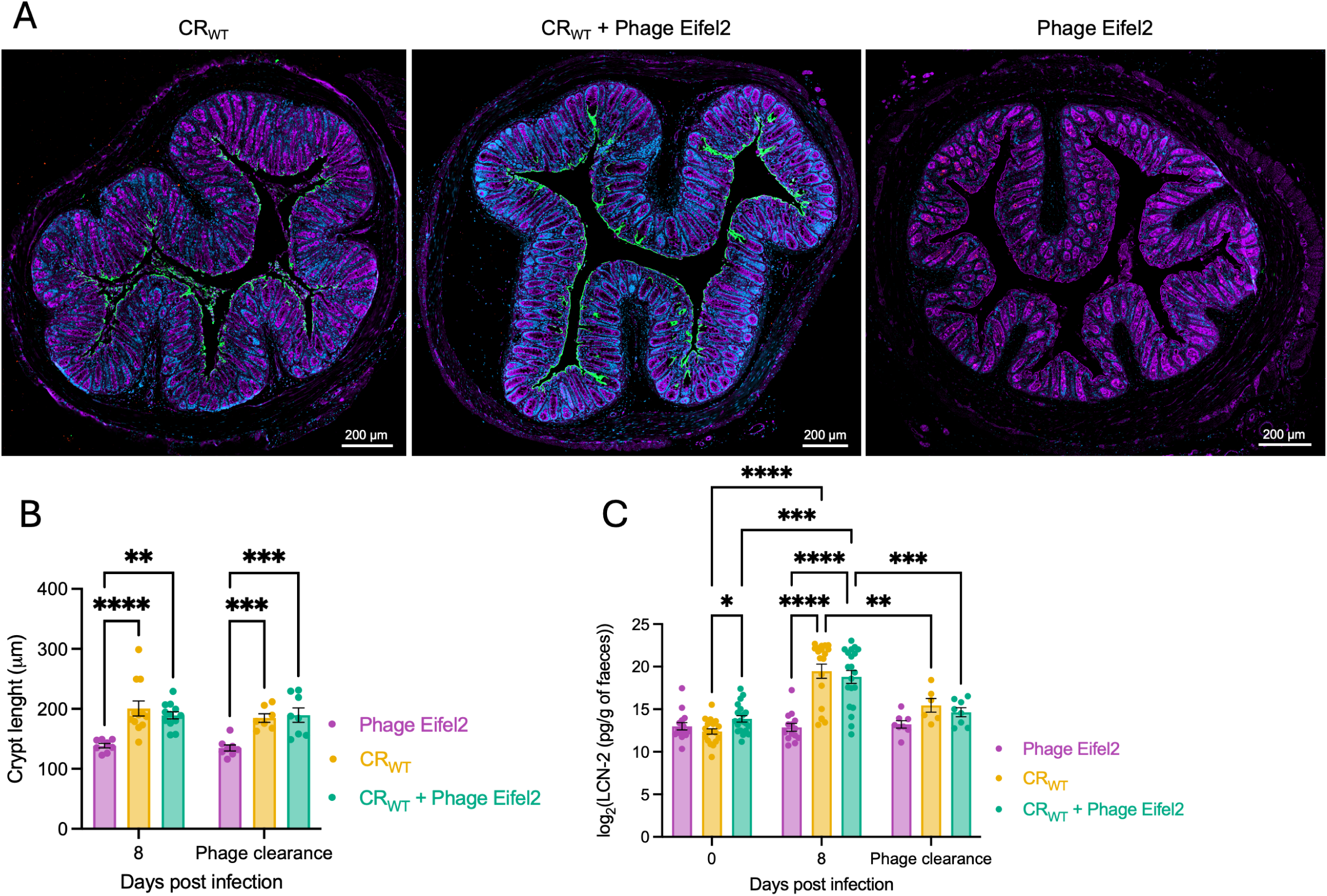
Eifel2 does not impact CR-induced colonic damage. (A) Representative immunofluorescence images of colonic sections at 8 dpi, with CR (green), anti-Eifel2 serum (red), WGA (purple) and DAPI (blue). Scale bars: 200 μm. (B) Crypt length measurements in distal colon sections at indicated timepoints. Data are shown as mean ± SEM and each dot represents the mean crypt length for an individual mouse. Statistical analysis was performed using two-way ANOVA with Tukey’s multiple comparison test. (C) LCN-2 levels in homogenised faeces at indicated timepoints, measured by ELISA. Data are shown as mean ± SEM and dots represent values for individual mice. Statistical analysis was performed using a mixed-effects model with Geisser-Greenhouse correction and Tukey’s multiple comparison test. (B, C) Statistical significance is indicated as: *P* < 0.05 (*); *P* < 0.01 (**); *P* < 0.001 (***); *P* < 0.0001 (****); non-significant comparisons are not shown.

To confirm these results, we quantified faecal levels of LCN-2, a marker of intestinal inflammation ^22^. At 8 dpi, LCN-2 levels were elevated in both infected groups, regardless of phage treatment, while phage treatment alone did not significantly alter this response (Fig. 5C). At the time of phage clearance, LCN-2 levels were comparable to pre-infection levels across all groups. These results suggest that oral administration of phage alone does not elicit an inflammatory immune response and that during stable Eifel2 – CR coexistence in the gut, the phage does not affect host – pathogen interactions.

### Emergence and characterisation of phage resistance *in vivo*

We next investigated the emergence of Eifel2-resistant CR *in vivo*. We screened bacterial isolates from mice faeces at 8 and 13-15 dpi using cross-streak assays. From each mouse, we tested up to six colonies for their susceptibility to Eifel2. No resistant colonies were detected in CR-infected, phage-untreated mice, while low levels of resistance were observed in the phage-treated infected group, with the percentage of resistant colonies ranging from 0% to 8.33% at 8 dpi (Table 1). At later timepoints, variability between mice increased, with the infected phage-treated group from one biological replicate reaching 63.64% resistant colonies, while others remained at 0%. Overall, resistant colonies were identified in only 6 of 20 mice that received phage treatment. Interestingly, in mice where resistant colonies were observed at 8 dpi, they were not detected at the later timepoint, suggesting that the selective pressure exerted by the presence of the phage was not enough to provide an advantage to the resistant clones, thus preventing the expected clonal expansion of the resistant phenotype in the gut. Thus, phage resistance seems disadvantageous in the context of coexistence. In agreement with these observations, the presence of resistant variants was not linked to slower CR clearance, thus confirming Eifel2 is not capable of driving CR clearance and selection of resistance clones at a population level.

**Table 1.**
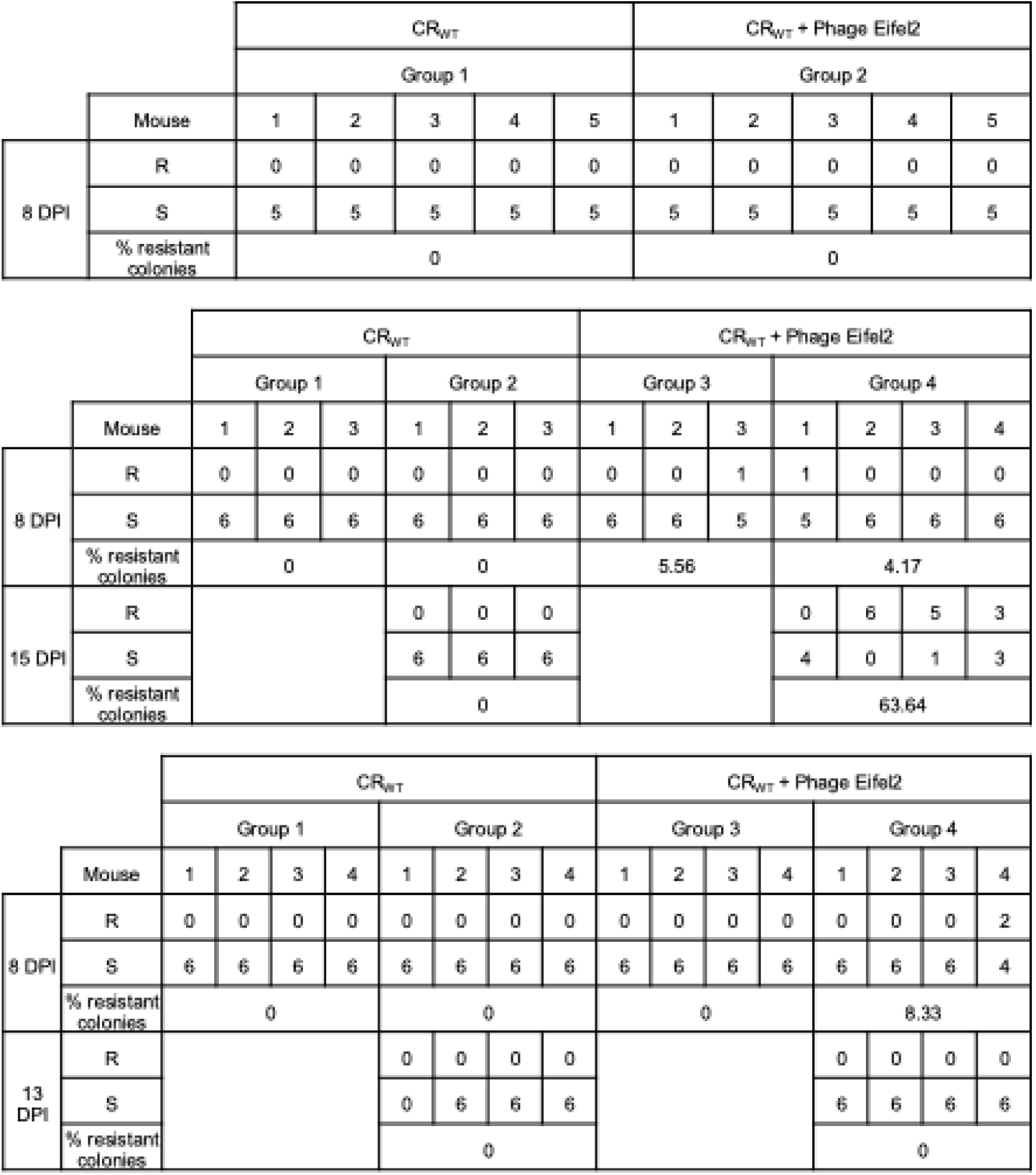
Quantification of resistant CR colonies in mouse faeces.

To investigate the genetic basis of resistance, one resistant colony from each of the six mice where resistance was observed (labelled R1 to R6) was sequenced and compared to the reference CR ICC168 genome (NCBI accession number: NC_013716.1) (Table S1). We identified three mutations that could be implicated in resistance: *rfaJ* and *rfaK*, both involved in lipopolysaccharide (LPS) biosynthesis (Fig. 6A); and *dsbC*, a gene encoding a periplasmic disulfide bond isomerase. A premature stop codon at position 307 of 1017 bp was identified in *rfaJ* in one isolate (R2), while transposon insertions were found at different positions in *rfaK* in two other isolates (R1 and R6). A large deletion affecting *dsbC* was detected in three isolates (R3, R4, and R5), with the deletion also disrupting the downstream gene *recJ*. None of the sequenced isolates exhibited a significant growth defect when grown in LB for 16 h (Fig. S5A).

**Figure 6.**
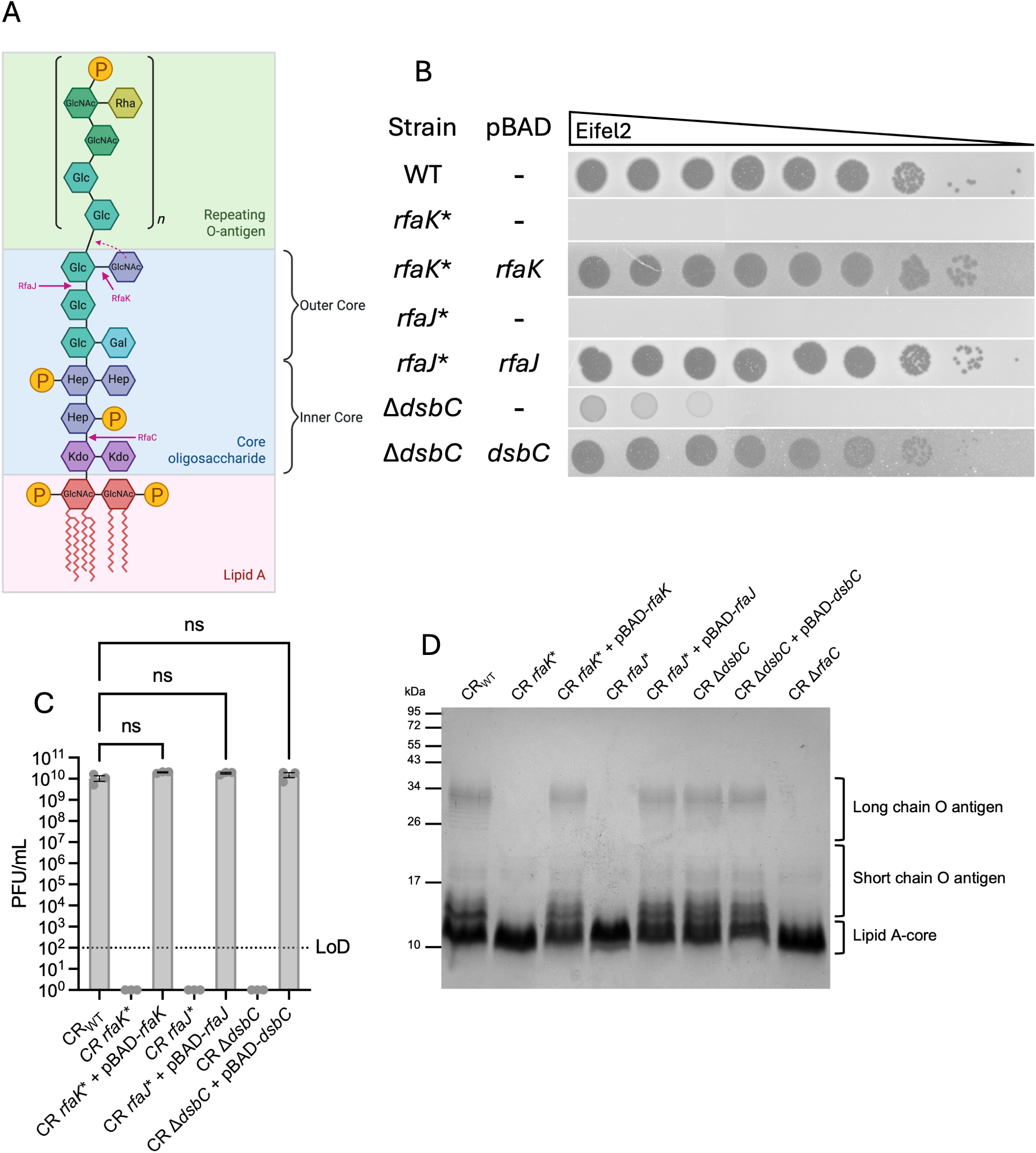
Genetic and structural basis of CR resistance to Eifel2. (A) Schematic representation of the CR LPS structure, highlighting the roles of RfaK, RfaJ and RfaC in structure assembly, inferred from structural homology with the *E. coli* type R2 *rfa* (*waa*) locus ^23–29^. GlcNAc: N-acetyl-D-glucosamine; P: phosphate; Kdo: 3-deoxy-D-manno-oct-2-ulosonic acid; Hep: heptose; Glc: D-glucose; Gal: D-galactose; Rha: L-rhamnose. (B-D) As *rfaJ* mutant could not be generated, the sequenced mutated faecal isolate (R2) was used, along with its complementation. (B) Representative images of DLA spot assays of Eifel2 on CR mutants and their complemented strains grown under pBAD-inducing conditions (0.2% L-arabinose). Images are representative of n = 3 independent biological replicates. (C) PFU counts of DLA spot assays of Eifel2 on CR mutants and their complemented strains grown under pBAD-inducing conditions. Values below the LoD were set to 1 for plotting. Data are shown as mean (SEM) of n = 3 independent biological replicates and dots represent PFU counts for individual repeats. Statistical analysis of strains with PFU counts above the LoD was performed using nonparametric Kruskal-Wallis test with Dunn’s multiple comparisons test. ns: non-significant. (D) Silver-stained SDS-PAGE gel of LPS extracted from each strain following 3 h of growth in pBAD-inducing conditions.

To study the spread of these mutations in our mouse model, we performed PCRs on the 18 isolated resistant colonies (including the 6 that were sequenced) to detect alterations in *rfaK*, *rfaJ* and the *dsbC*-*recJ* region (Fig. S5B). In the case of *rfaK*, three colonies isolated from two different experiments produced fragments consistent with the insertion of a transposon. For *dsbC* and *recJ*, 14 colonies isolated from the same cage produced PCR products consistent with the deletion observed in the sequenced isolates.

To confirm the role of these mutations in resistance, we generated clean mutants in the wild-type CR background. A stop codon mutation was introduced in *rfaK*, and a deletion mutant was generated for *dsbC*. Because the deletion affecting *dsbC* also disrupted the downstream gene *recJ* in the sequenced isolate, we also generated a stop codon mutant in *recJ*. Plating assays showed that mutations in *rfaK* and *dsbC* conferred full resistance to Eifel2 infection (Fig. 6B, C). The *dsbC* mutant showed low levels of lysis at high phage concentrations but did not support plaque formation (Fig. 6B,C), similar to the phenotype observed in the sequenced isolates (Fig. S5C). Complementation of the mutants with wild-type *rfaK* or *dsbC* restored full susceptibility, confirming the role of these mutations in resistance. As attempts to generate mutants for *rfaJ* were unsuccessful, we complemented the original resistant R2 isolate with a plasmid carrying the wild-type *rfaJ* gene instead. Susceptibility of the complemented R2 isolate to Eifel2 infection was fully restored, confirming the role of the mutation in *rfaJ* in resistance (Fig. 6B, C). In contrast, the *recJ* mutation did not affect phage susceptibility (Fig. S5D). Plating assays with the unrelated phage ColRes showed a slightly lower plating efficiency with *rfaK* and *rfaJ*, but not *dsbC* mutants (Fig. S5E), although no strain showed efficient resistance to ColRes infection, indicating the resistance mechanisms are specific to Eifel2 infection.

The involvement of *rfaJ* and *rfaK* in core LPS biosynthesis points towards a component of LPS being the receptor for the phage. We examined LPS profiles of mutant and complemented strains by silver staining. We used the wild-type strain and a known deep-rough mutant (Δ*rfaC*), which was also resistant to Eifel2 (Fig. S5F), as controls. Both *rfaJ* and *rfaK* mutants showed truncated LPS with a lack of O-antigen, agreeing with the presence of the terminal GlcNAc, added by RfaK, being essential for O-antigen ligation ^23,29^, whereas the complemented strains restored normal LPS length (Fig. 6D). The *dsbC* deletion mutant did not show any noticeable differences in LPS structure, suggesting that resistance is mediated by a different mechanism. To rule out O-antigen component of the LPS as receptor for Eifel2, a CR strain lacking O-antigen was tested. The O-antigen deficient mutant remained sensitive to Eifel2 (Fig. S5F), suggesting that while the *rfaJ* and *rfaK* mutants lack O-antigen, the resistance is due to the mutation affecting the core LPS components.

## Discussion

CR has extensively been used as a translational model to study EPEC and EHEC infections *in vivo*, given its ability to infect immunocompetent mice in the presence of the natural microbiota ^12–14^. Although CR-specific phages have been isolated, their use *in vivo* remains unexplored ^15–18^. Thus, the aim of this study was to isolate lytic phages infecting CR and develop an *in vivo* model of phage therapy to accurately depict phage-pathogen interactions in a physiological context.

We isolated two closely related lytic phages, Eifel1 and Eifel2, infecting CR. Genomic analysis revealed they represent a new species within the *Felixounavirus* genus. Phages within the *Felixounavirus* genus are strictly lytic and primarily target Gram-negative enterobacteria, including *Salmonella* spp. and *E. coli*, and their potential as antimicrobial agents has been previously explored *in vitro* ^30,31^. Despite almost identical genomes, Eifel2 exhibited higher lytic activity compared to Eifel1, potentially due to a structural variation in a minor tail protein, which could enhance host attachment and/or DNA injection efficiency ^32,33^. An additional gene in Eifel2, located among hypothetical proteins, remains functionally uncharacterised and needs further investigations to explore its potential implications in the infection cycle of Eifel2. Interestingly, both phages displayed strict host specificity for CR, in contrast to other members of the *Felixounavirus* genus ^34^.

To visualise phage infection dynamics, we developed a simplified immunisation protocol using whole phage particles. Following two rounds of immunisation, the resulting serum recognised multiple virion proteins by Western blot. Compared to previous approaches such as fusion of fluorescent tags to capsid proteins ^35,36^, fluorescent labelling of virions ^37,38^ or phage DNA ^39–42^, FISH-based phage DNA detection ^43,44^, plasmid-based approaches ^45^, and generation of capsid protein specific antibodies ^46^, our method provides key advantages: no genetic manipulation of phages, avoiding potential disruptions to phage infectivity; visualisation of multiple infection cycles and phage progeny, not limited to initial input phages; applicability to unsequenced phages, enabling broader use and broad antigen detection and bypassing the need to purify individual phage proteins.

In the CR mouse infection model, Eifel2 showed robust replication in the gut. However, phage treatment did not significantly reduce bacterial burden or alleviate infection-associated colonic pathology. A strong positive correlation between phage and bacterial titres was observed, suggesting establishment of dynamic coexistence rather that phage-mediated bacterial clearance. This is consistent with previous *in vivo* studies in murine models with different *E. coli* strains, that showed stable coexistence of phages and host bacteria in the gut ^5,7,47^. Those studies relied on disturbing the microbiome to allow for pathogen colonisation, either by antibiotic pre-treatment to use enteroaggregative O104:H4 *Escherichia coli* (EAEC) ^9,50^, or using mice with a synthetic microbiota like the OMM^12^ mouse model, to infect with *E. coli* Mt1B1 and EAEC^11^. Notably, Lourenço et al. ^7^ showed that coexistence was enabled by the spatial partitioning of bacteria and phages in the gut: phage amplification occurred mainly in the lumen, thus, bacteria adhering to mucosal surfaces were protected from phage predation. Upon infection, CR intimately attaches to intestinal epithelial cells (IECs) and are shed into the lumen ^48^, and could thus support this model of niche partitioning.

Consistently, immunofluorescence imaging using the generated serum failed to detect phage signal in distal colon sections, despite high faecal titres. This could reflect low phage abundance at the mucosal surface in the colon, consistent with the hypothesis of phage replication occurring in the lumen and does not exclude the application of the generated serum in further *in vivo* applications.

A recent study using *E. coli* Mt1B1-colonised OMM^12^ mice showed that emergence of phage resistance is not a major contributor to phage-bacteria coexistence in the gut ^7^. In our model, phage-resistant mutants emerged at low frequencies. However, these mutants did not clonally expand and spread, suggesting a significant fitness cost for phage resistance in the highly selective gut environment ^1,2^. Spatial segregation of phages and bacteria in the gut could further explain the low emergence frequencies of resistant isolates by limiting selection ^7^. Whole genome sequencing of resistant isolates revealed resistance-conferring mutations in *rfaJ* and *rfaK*, genes involved in core LPS biosynthesis ^49^, that resulted in truncated LPS and loss of O-antigen. These mutations have previously been described in phage resistance mechanisms in Gram-negative bacteria ^50–52^. However, O-antigen is an important virulence factor for *in vivo* colonisation and immune evasion in Gram-negative pathogens ^53^, potentially explaining the low frequency and limited expansion of these mutants. A large deletion affecting *dsbC*, a gene encoding a periplasmatic disulfide bond isomerase ^54–57^, was observed more frequently. *dsbC* mutants exhibited intact LPS and O-antigen, possibly contributing to a lower fitness cost *in vivo*, and thus a more important spread of this mutation between mice within a cage. However, these mutants also displayed partial sensitivity to high concentrations of phage lysate without supporting plaque formation.

Absence of DsbC indirectly affects bacterial resistance to environmental elements, such as oxidative stress ^58^, suggesting CR *dsbC* mutants could exhibit lower resistance to certain elements present in phage lysate.

The precise mechanisms by which core LPS and *dsbC* mutants confer resistance to Eifel2 remain unclear but seem to be highly specific. Given DsbC’s role in protein folding, loss of function could lead to alterations in membrane proteins or alter membrane composition, potentially impairing phage receptor availability. As O-antigen deficient strains remained sensitive, major LPS core modifications, such as the ones observed in *rfaK* and *rfaJ* mutants, could lead to similar effects.

Taken together, this study establishes a murine model to investigate phage-bacteria interactions *in vivo* in the context of the complete endogenous gut microbiota. Our results showed that despite strong *in vitro* lytic activity, oral administration of Eifel2 resulted in the emergence of a dynamic and spatially structured system that supports coexistence rather than eradication of one or the other in the gut environment. We have shown that while resistance emerged, the resistant CR mutants did not clonally expand, indicating an associated fitness-cost. Our findings highlight the importance of studying phage-bacteria interactions in physiologically relevant models for revealing the true complexity of such interactions.

## Materials and methods

### Bacterial strains and growth conditions

All bacterial strains used in this study are listed in Table S2. Strains were cultured in Lysogeny broth (LB) at 37 °C with shaking at 200 rpm, or on LB agar (1.5%) plates incubated statically at 37 °C. Where appropriate, antibiotics were added to the media at the following final concentrations: nalidixic acid (Nal), 50 μg/mL; gentamicin (Gm), 10 μg/mL; kanamycin (Km), 50 μg/mL; and streptomycin (Sm), 50 μg/mL.

For experiments involving complemented strains harbouring the pBAD expression plasmid, cultures were grown in LB supplemented with 0.2% w/v glucose to repress expression. To induce gene expression, 0.2% w/v L-arabinose was added to the media in place of glucose.

### Generation of CR mutants and complementation

Plasmids and primers used in this study are listed in Table S2. All genomic deletions and point mutations were generated using a two-step recombination protocol, resulting in scarless and markerless mutants, as previously described by ^59^. Genomic DNA (gDNA) from CR ICC169 was extracted using the Monarch Genomic DNA Purification Kit (New England Biolabs, NEB), following manufacturer’s instructions. For gene deletions, ∼500 bp homology regions (HRs) flanking the gene of interest were PCR amplified using 2X Phanta Flash Master Mix (Vazyme) from CR gDNA. To generate point mutations, ∼500 bp HRs flanking the mutation site were amplified using primers incorporating a stop codon. PCR products were purified using the Monarch PCR & DNA Cleanup Kit (NEB). The pSEVA612S_revISceI vector was linearised via PCR using 2X Phanta Flash Master Mix (Vazyme) and purified using the same cleanup kit. HRs were inserted into the linearised vector using Gibson Assembly (NEB), following manufacturer’s instructions. Assembled vectors were transformed into chemically competent *E. coli* CC118λpir cells and maintained with appropriate antibiotics.

To introduce the desired mutations into CR, recipient strains pre-transformed with pACBSR (plasmid expressing the I-SceI endonuclease and lambda red fragment) were used. Tri-parental conjugation was performed by incubating 20 μL of the donor strain (*E. coli* CC118λpir carrying the mutagenesis vector) with 20 μL of the helper strain (*E. coli* 1047 carrying pRK2013) on LB agar at 37 °C for 2 h. Then, 40 μL of the recipient strain (CR pACBSR) was added and incubated for at least 8 h at 37 °C. Transconjugants were selected on LB agar plates containing appropriate antibiotics. Growing colonies were grown in LB supplemented with L-arabinose (0.4%) and appropriate antibiotics for 2 h to induce I-SceI expression. Cultures were plated on LB agar with appropriate antibiotics, and candidate deletion mutants were screened via colony PCR using 2X Rapid Taq Master Mix (Vazyme).

For genetic complementation, genes of interest were PCR-amplified from CR gDNA and the pBAD vector was linearised by PCR using 2X Phanta Flash Master Mix (Vazyme). PCR amplicons were purified using Monarch PCR & DNA Cleanup Kit (NEB), and genes were inserted using Gibson Assembly (NEB). Resulting plasmids were transformed into electrocompetent mutant CR pACBSR cells prepared at room temperature as previously described ^60^ and selected on LB agar supplemented with glucose (0.2%) and appropriate antibiotics. For *rfaJ* complementation, the assembled plasmid was introduced into the sequenced faecal bacterial isolate and selection was done on LB agar with glucose (0.2%) and appropriate antibiotics. All plasmids and mutants were validated by colony PCR using 2X Rapid Taq Master Mix (Vazyme) and Sanger sequencing (Eurofins Genomics).

### Phage isolation and purification

Double layer agar (DLA) assays were performed as previously described ^61^, with minor modifications. Briefly, 4 mL of soft LB agar (0.3%) supplemented with 10 mM MgSO₄ and 10 mM CaCl₂ were inoculated with 100 μL of overnight culture of the bacterial host. For DLA plate assays, the inoculated soft agar was mixed with 100 μL of phage suspension before being poured onto LB agar (1.5%) plates supplemented with appropriate antibiotics in standard 90 mm Petri dishes. For DLA spot assays, only the bacterial culture was mixed into the soft agar, poured onto LB agar (1.5%) plates, and allowed to solidify before spotting 10 μL of serially diluted phage suspensions onto the surface. Plates were incubated overnight at 37 °C under static conditions. For larger square Petri dish plates, reagent volumes were doubled.

All phages isolated in this study are listed in Table S2. Sewage water samples were initially centrifuged and filtered through 0.45 μm pore-size membranes. The filtrate was added to LB supplemented with Nal inoculated with CR and incubated overnight at 37 °C with shaking. The resulting co-cultures were centrifuged and filtered through 0.45 μm membranes to obtain crude phage lysates, which were serially diluted, and DLA plate assays were performed. For single plaque isolation and phage purification, a prophage-free CR strain (CR Δ10, a derivative of ICC169) was used in all subsequent steps. Single plaques were picked and placed into saline-magnesium (SM) buffer (100 mM NaCl, 8 mM MgSO₄, 50 mM Tris-HCl pH 7.5) for 2 h at room temperature. This purification process was repeated three times.

High-titre phage lysates were generated either by liquid co-culture (Eifel2) or large-scale DLA plate assays (Eifel1 and ColRes). For the latter, ∼20 DLA plates were inoculated with appropriate phage dilutions to achieve near-confluent lysis (“webbed” plates). The following day, 5 mL of SM buffer were added to each plate and incubated with gentle shaking for 2 h at room temperature. The buffer was then collected, pooled, centrifuged, and filtered through 0.45 μm membranes. High-titre lysates were used to inoculate large liquid co-cultures to obtain large high-titre phage preparations for downstream purification steps. These lysates were concentrated by ultracentrifugation and purified by CsCl density gradient ultracentrifugation. Briefly, pooled lysates were centrifuged at 40,000 × g for 2 h, and the supernatant was discarded. This step was repeated as needed to concentrate phage particles. The resulting pellets were resuspended in 1 mL SM buffer and incubated overnight at 4 °C, before being pooled. CsCl gradients were prepared with layers of 1.7, 1.5, and 1.3 g/cm³ CsCl solutions in SM buffer. Gradients were ultracentrifuged at 40,000 rpm for 3 h at 5 °C. Visible phage bands were extracted using a sterile needle and syringe and subsequently washed and concentrated using centrifugal concentrators (100,000 kDa MWCO, Vivaspin) with sterile SM buffer according to manufacturer’s instructions. Final phage preparations were filtered with 0.45 μm membranes and were stored in SM buffer at 4 °C.

### TEM images

TEM of phage preparations was performed by applying 2-3 μL of purified phage to glow-discharged, carbon-coated copper grids (200 mesh, Agar Scientific), and then staining with 2% uranyl acetate. Micrographs of phages were captured on an FEI T12 electron microscope at an acceleration voltage of 120 kV and a nominal magnification of 26,000x.

Images were processed and analysed using Zen 3.5 Blue software (Carl Zeiss MicroImaging GmbH, Germany). Phage dimensions were determined by measuring at least 20 individual virions. Measurements included capsid length and width, tail length and neck length, and the mean values were calculated for each parameter.

### Phage lysis curves

Overnight cultures of CR_WT_ were diluted in LB supplemented with Nal to reach an initial OD₆₀₀ of 0.1. Dilutions of purified phage suspensions were prepared in SM buffer and added to the diluted CR cultures to achieve the indicated MOIs. Control wells received SM buffer without phage. Each culture was loaded into a 96-well flat-bottom microplate in technical triplicates, and OD₆₀₀ readings were recorded every 15 min over 16 h using a FLUOstar Omega microplate reader (BMG Biotech) at 37 °C with continuous shaking.

### Bacterial growth curves

Overnight cultures of bacterial strains were diluted in LB supplemented with appropriate antibiotics to reach an initial OD₆₀₀ of 0.1. Cultures were aliquoted into a 96-well flat-bottom microplate in technical duplicates and OD₆₀₀ measurements were recorded every 15 min for 16 h using a FLUOstar Omega microplate reader (BMG Biotech) at 37 °C with continuous shaking.

### One-step growth curve assay

One-step growth curves were performed using a modified protocol based on ^62^. Briefly, CR cultures were grown to mid-log phase (OD₆₀₀ = 0.2) and infected with purified phage suspensions at a MOI of 1. Cultures were incubated at 37 °C with shaking for 5 min to allow phage adsorption. Following adsorption, cultures were centrifuged to remove unbound phages, and pellets were resuspended in fresh LB supplemented Nal. The infected cultures were then incubated at 37 °C with shaking. Samples were collected at the indicated timepoints, immediately serially diluted in SM buffer, and phage titres were determined using DLA spot assays.

### Phage DNA extraction and sequencing

Phage DNA was extracted using the MasterPure™ Complete DNA and RNA Purification Kit (Lucigen) following an optimised protocol adapted from the “DNA Purification Protocols – Fluid Samples” section of the manufacturer’s manual. Briefly, 150 μL of high-titre filter-sterilised purified phage lysate was mixed with 150 μL of 2X Tissue and Cell Lysis Solution supplemented with 1 μL RNase A (5 μg/mL) and 1 μL RNase-free DNase I (1 U/μL). Samples were incubated at 37 °C for 30 min to remove contaminating DNA and RNA. To inactivate nucleases, 20 μL of 0.5 M EDTA was added, followed by 1 μL of Proteinase K, and samples were incubated at 65 °C for 15 min, vortexing every 5 min. Samples were then cooled on ice for 5 min before the addition of 150 μL of MPC Protein Precipitation Reagent. Tubes were vortexed and centrifuged at 4 °C. The supernatant was transferred to a clean microcentrifuge tube and mixed with 500 μL of isopropanol. DNA was pelleted by centrifugation at 4 °C, and the isopropanol was carefully decanted. Pellets were washed twice with 70% ethanol, ensuring not to dislodge the pellet, and residual ethanol was removed completely. Purified DNA was resuspended in 15 μL of Elution Buffer (EB) (Qiagen) and submitted for whole genome sequencing through the Enhanced Genome Service provided by MicrobesNG, following their standard submission protocols. Sequencing was performed using a combination of Illumina short-read and Oxford Nanopore long-read technologies.

### Assembly, annotation, visualisation and alignment of phage genomes

Prior to genome assembly, host-derived (*Citrobacter rodentium* ICC168, NC_013716.1) sequence contamination was removed from raw reads using bbduk (BBTools v39.13). Decontaminated reads were assembled using SPAdes genome assembler ^63,64^ (v4.0.0, hybrid assembly using --nanopore option and --isolate options, default options were used for the other parameters) and annotation of assembled contigs was performed using Pharokka ^65^ (v1.7.4, default options with the addition of --dnaapler to reorder the genome on the *terL* gene) and Phold (v0.2.0, using the run module with the --ultra-sensitivity option) (https://github.com/gbouras13/phold). Assemblies and annotations were visualised and curated using Geneious Prime® 2025.1.2 (GraphPad Software LLC d.b.a Geneious).

Taxonomic classification of phages genomes was carried out using the web-based taxMyPhage tool ^66^. Phylogenetic relationships between species within the same genus were inferred using VICTOR and visualised as taxonomic trees. The entire analysis was carried out by the VICTOR web service (https://victor.dsmz.de), a method for the genome-based phylogeny and classification of prokaryotic viruses ^67^. All pairwise comparisons of the nucleotide sequences were conducted using the Genome-BLAST Distance Phylogeny (GBDP) method ^68^ under settings recommended for prokaryotic viruses ^67^. The resulting intergenomic distances were used to infer a balanced minimum evolution tree with branch support via FASTME including SPR postprocessing ^69^ for each of the formulas D0, D4 and D6, respectively. Branch support was inferred from 100 pseudo-bootstrap replicates each. Trees were rooted at the midpoint ^70^ and visualized with ggtree ^71^. Taxon boundaries at the species, genus and family level were estimated with the OPTSIL program ^72^, the recommended clustering thresholds ^67^ and an F value (fraction of links required for cluster fusion) of 0.5 ^68^. The online PhageAI tool ^73^ was used to predict the lifestyle of phages.

Whole genome alignments were performed with the progressive Mauve algorithm ^19^ and visualised in Geneious Prime® 2025.1.2 (GraphPad Software LLC d.b.a Geneious), and whole genome comparisons were visualised using Easyfig ^20^. Protein structural predictions were performed using the web version of AlphaFold 3 ^74^. Resulting models were visualised and analysed using UCSF ChimeraX (v1.9) ^75^.

Assembled genomes are available at https://doi.org/10.5281/zenodo.15724312.

### Phage pH and temperature stability

To assess temperature stability, purified phage suspensions in SM buffer were incubated for 24 hours at the indicated temperatures. At designated time points, samples were collected and PFUs were determined using a DLA spot assay. To assess phage stability upon freezing, multiple aliquots of the same phage suspension were stored at –20 °C to avoid repeated thawing and refreezing. PFUs were determined after thawing a single-use aliquot. To assess pH stability, purified phage suspensions in SM buffer were diluted 1:100 in SM buffer adjusted to the indicated pH values and incubated at 37 °C for 24 hours in a static incubator. Following incubation, PFUs were quantified using DLA spot assays.

### Mouse experiments

Mouse experiments were performed in accordance with the Animals Scientific Procedures Act of 1986 ^76^ and UK Home Office guidelines and were approved by the Imperial College Animal Welfare and Ethical Review Body. Experiments were designed in agreement with the ARRIVE guidelines ^77^ for the reporting and execution of animal experiments. Licence number PPL PP7392693.

Specific pathogen-free C57BL/6 female mice (18-20 g; 6-8 weeks) and pathogen-free CD-1 female mice (29-31 g; 5-7 weeks) were purchased from Charles River Laboratories and housed in groups of 2 to 5 individuals in high-efficiency particulate air (HEPA)-filtered cages with bedding, nesting and free access to food and water. The temperature, humidity, and light cycles were kept within the UK Home Office code of practice, with the temperature between 20°C and 24°C, the room humidity at 45-65 %, and a 12 h/12 h light cycle with a 30-min dawn and dusk period to provide a gradual change. We used female mice only to avoid the logistical issues and stress to animals associated with male in-cage fighting. For each experiment, mice were randomly assigned to experimental groups. Investigators were not blind to the allocation. Treatments and measurements were always performed following the same cage order.

### Mouse infections and phage treatment

ICC169 (CR_WT_) was cultured overnight in LB supplemented with Nal. The following day, saturated bacterial cultures were centrifuged, and pellets were resuspended in 1.5 mL of sterile phosphate-buffered saline (PBS). C57BL/6 mice were infected by oral gavage with 200 μL of the bacterial suspension (∼1 × 10⁹ CFUs), or mock-infected with 200 μL of sterile PBS. The inoculum dose was retrospectively confirmed by serial dilution and CFU enumeration on LB agar supplemented with Nal.

On days 3, 4, 5, 6, and 7 post-infection, mice received 200 μL of a sterile purified phage Eifel2 suspension (1 × 10⁹ PFUs) in SM buffer by oral gavage. Control animals received 200 μL of sterile SM buffer. To prevent phage adsorption to mouse feed and increase stomach pH, thus enhancing phage treatment efficacy, food was withdrawn 3 h prior to phage administration and reinstated immediately afterward. Body weight and bacterial burden in faeces were monitored throughout the infection as previously described ^14^. Briefly, faecal pellets were collected on indicated days, weighed, and homogenised in PBS (0.1 g of stool/mL of PBS) using a vortex mixer for 15 min at room temperature. Homogenates were serially diluted and plated on LB agar supplemented with Nal to determine CFU counts. For phage quantification, homogenised faecal samples were centrifuged and filtered through 0.45 μm pore-size membranes. Filtrates were serially diluted and spotted using a DLA-spot assay as described above.

At 8 dpi, mice were culled by cervical dislocation. At 21 or 23 dpi, mice were weighed and anesthetised with ketamine (100 mg/kg) and medetomidine (1 mg/kg) administered via intraperitoneal injection using a 27G, 13 mm needle (BD microlance). Once mice had lost pedal reflexes, whole blood was collected by cardiac puncture (25G, 16 mm needles, BD microlance) and transferred to a serum collection tube containing a clot activator and separating gel (BD Microtainer SST, BD). Mice were then euthanised by cervical dislocation and caecal contents and distal colon sections were harvested for downstream analyses. CFU and PFU counts in caecal contents were determined using the same procedures described above for faecal samples.

### Mice Immunisation

CD-1 mice were immunised via subcutaneous injection with 2.2 × 10⁸ PFUs of sterile, purified Eifel2 phage particles in 220 μL of SM buffer, supplemented with 30 μL of alum adjuvant (Alum Alhydrogel® adjuvant 2%). Control mice received 220 μL of sterile SM buffer with 30 μL of alum. A booster injection was administered 21 days after the initial immunisation. Body weight loss of mice was monitored throughout the experiment. At 21 days post-booster injection, ∼30 μL of blood was collected via tail vein sampling and allowed to clot at room temperature to assess serum reactivity to Eifel2. At 22 days post-booster injection, mice were first anesthetised as described above. Whole blood was then collected by cardiac puncture as above for subsequent serum processing and use.

### Isolation of resistant colonies from faecal samples and phage host range determination

Resistance of faecal bacterial isolates and the host range of isolated phages were assessed using cross-streak assays. Colonies that grew from plated faecal samples were picked and cultured overnight in LB supplemented with Nal at 37 °C with shaking. Bacterial strains for host range determination were cultured overnight in LB at 37 °C with shaking. To perform cross-streak assays, 50 μL of purified high-titre phage suspension was streaked in a single horizontal line across a LB agar square plate and allowed to dry. Then, 10 μL of overnight bacterial cultures was streaked perpendicularly to the phage streak. Plates were incubated overnight at 37 °C under static conditions. The following day, bacterial lysis at the intersection of the bacterial and phage streaks was recorded as an indicator of phage sensitivity.

### Whole genome sequencing of resistant isolates and mutation identification

Resistant bacterial colonies isolated from faecal samples were submitted for whole genome sequencing through the Enhanced Genome Service provided by MicrobesNG, following their standard submission protocols. Sequencing was performed using a combination of Illumina short-read and Oxford Nanopore long-read technologies.

SNPs were identified using Snippy (https://github.com/tseemann/snippy), while larger genomic rearrangements and insertions/deletions were detected using Pilon ^78^. Both analyses were carried out using the Galaxy Europe platform (https://usegalaxy.eu) ^79^.

To validate mutations or assess integrity of target genes across all isolated resistant colonies, PCR amplification of the *rfaK*, *rfaJ*, and *dsbC/recJ* regions was performed using 2X Rapid Taq Master Mix (Vazyme) with appropriate primers annealing immediately upstream or downstream of each gene or region. Amplified products were resolved on 1% agarose Tris-Acetate-EDTA (TAE) gels containing SYBR Safe DNA Gel Stain (1:10,000) and electrophoresed under standard conditions. Gels were imaged using a Chemidoc imaging system (Bio-Rad) and analysed using ImageLab (v6.1) (Bio-Rad).

### LCN-2 ELISA

Mouse faecal pellets were weighed and homogenised in PBS (0.1 g of stool/1 mL of PBS) using a vortex mixer for 15 min at room temperature. Samples were centrifuged to remove debris, and the supernatants were collected and stored at −80 °C. Faecal LCN-2 concentrations were measured using a DuoSet Mouse Lipocalin-2 ELISA kit (Bio-Techne), following manufacturer’s instructions. Absorbance readings were taken with a FLUOstar Omega microplate reader (BMG Labtech).

### Serum collection and processing, and anti-Eifel2 IgG ELISA

Mice were anesthetised and blood was collected via cardiac puncture as described above. Blood was left to clot for 1 h at room temperature, serum was subsequently separated by centrifugation at 20 000 x g for 3 mins and stored in aliquots at −80 °C until use or further analysis.

For quantification of anti-Eifel2 IgG levels in serum, ELISA clear flat-bottom Maxisorp immune 96-well plates (Thermo Fisher) were coated overnight at 4 °C with either 100 μL/well of purified Eifel2 phage suspension (1 × 10^10^ PFU/mL) for sample wells or 100 μL/well of anti-mouse IgG antibody (2 μg/mL) for standard wells. Plates were washed three times with PBS containing 0.1% Tween 20 (PBS-T), then blocked for 1 h at room temperature with PBS-T supplemented with 5% bovine serum albumin (BSA). Serum samples and serial dilutions of purified mouse IgG as the standard were added to the appropriate wells and incubated overnight at 4 °C. Plates were washed as described above and incubated with horseradish peroxidase (HRP)-conjugated goat anti-mouse IgG antibody (1:2000; Jackson ImmunoResearch) for 2 h at room temperature. Plates were washed as described above, and detection was carried out by adding 100 μL/well of 3,3′,5,5′-tetramethylbenzidine (TMB) substrate. The reaction was stopped after 5–10 min with 2N orthophosphoric acid.

Absorbance was measured at 450 nm and at 540 nm using a FLUOstar Omega microplate reader (BMG Labtech). Anti-Eifel2 IgG concentrations were determined by comparison to the IgG standard curve.

### Western Blot and Dot Blot

For Western blotting, purified high-titre Eifel2 lysate (10^11^ PFU/mL) was mixed with 5× SDS loading buffer containing 0.1 M dithiothreitol (DTT), boiled at 95 °C for 5 min, vortexed, then boiled again at 95 °C for 5 min and vortexed again before centrifugation. Samples were resolved on a 12 % SDS-PAGE gel under standard electrophoresis conditions and transferred onto pre-activated PVDF membranes using a semi-dry transfer system (Bio-Rad). Membranes were blocked for at least 4 h at room temperature in 10% skimmed milk in PBS-T, then incubated overnight at 4 °C with polyclonal anti-Eifel2 mouse serum (1:500) in 5% milk PBS-T. Following incubation, membranes were washed three times (10 min each) in PBS-T and incubated for 1 h at room temperature with HRP-conjugated goat anti-mouse IgG antibody (1:5000) in 5% milk PBS-T. After three additional PBS-T washes (10 min each), membranes were developed using ECL substrate (Bio-Rad).

For dot blotting, 5 μL of purified phage suspensions (1 × 10⁹ PFU/mL) or 5 μL of filtered (0.45 μm membrane) CR_WT_ overnight culture supernatant were spotted onto dry nitrocellulose membranes. Once dry, membranes were blocked and processed identically to Western blot membranes, using the same antibody concentrations and incubation conditions. Blots were imaged using a Chemidoc imaging system (Bio-Rad) and were analysed using ImageLab (v6.1) (Bio-Rad).

### Phage *in vitro* staining

1 mL of CR Δ10 strain constitutively expressing GFP overnight culture was harvested by centrifugation, and the bacterial pellet was resuspended in 1 mL of purified high-titre Eifel2 lysate (10¹⁰ PFU/ml). Samples were incubated for 5 min at 37 °C with shaking to allow phage binding. Cells were pelleted again and resuspended in PBS with 1% paraformaldehyde (PFA) for 30 min on ice to fix. Following fixation, cells were washed twice with PBS and blocked by resuspension in PBS with 3% BSA for 1 h at room temperature with continuous agitation. After blocking, samples were pelleted and resuspended in the generated polyclonal anti-Eifel2 mouse serum (1:50 dilution in PBS with 1% BSA) and incubated overnight at 4 °C with continuous agitation. Cells were then washed twice in PBS and incubated with the secondary antibody (1:500) and DAPI (1:1000) (time course infection only) in PBS with 1% BSA for 1 h at room temperature in the dark with continuous agitation. After two final PBS washes, the final cell pellet was resuspended in 50 µl of PBS and mounted on microscopy slides using agarose pads. Details about the reagents and antibodies used can be found in Table S2.

For time-course imaging of phage infection, CR Δ10 strain constitutively expressing GFP were infected with phage Eifel2 following the protocol described above for one-step growth curve experiments (One-step growth curve assay section). At defined time points post-infection, samples were collected and processed for imaging using the *in vitro* staining protocol described above. Images were acquired using a Zeiss AxioVision Z1 microscope with a Hamamatsu digital camera. Images were processed and analysed using Zen (v3.5) Blue software (Carl Zeiss MicroImaging GmbH, Germany).

### Histological analysis and immunofluorescence staining

The distal 0.5 cm of the colon was harvested and fixed in 4% paraformaldehyde (PFA) for 2 h at room temperature, then transferred to 70% ethanol for storage. Tissues were processed, paraffin-embedded, and sectioned at 5 μm. Sections were stained with either hematoxylin and eosin (H&E) or processed for immunofluorescence. Crypt hyperplasia was quantified by measuring the lengths of at least 20 well-oriented colonic crypts per mouse from H&E-stained sections. The mean crypt length of each mouse was used for statistical analysis.

For immunofluorescence staining, paraffin-embedded sections were dewaxed and rehydrated by sequential submersion in Histo-Clear solution (2 × 10 min), 100% ethanol (2 × 10 min), 95% ethanol (2 × 10 min), 80% ethanol (2 × 3 min), and PBS containing 0.1% Tween 20 and 0.1% saponin (PBS-TS; 2 × 3 min). Antigen retrieval was performed by heating the slides for 30 min in demasking solution (0.3% trisodium citrate and 0.05% Tween 20 in distilled water, pH 6.0). Slides were cooled and then blocked in PBS-TS supplemented with 10% normal donkey serum (NDS) for at least 6 h in a humid chamber at room temperature. Sections were incubated overnight at 4 °C with primary antibodies diluted in PBS-TS containing 10% NDS. The following primary antibodies were used: rabbit polyclonal anti-CR serum (1:50) and mouse polyclonal anti-Eifel2 serum generated in this study (1:10). Slides were washed twice in PBS-TS (10 min each), followed by incubation with secondary antibodies (1:100), fluorescent wheat germ agglutinin (WGA) (1:200) and DAPI (1:1000). After two final PBS-TS washes (10 min each), slides were mounted using ProLong™ Gold Antifade Mountant (Thermo Fisher Scientific) and allowed to dry in the dark. Details about the reagents and antibodies used can be found in Table S2.

Images were acquired using a Zeiss AxioVision Z1 microscope. Fluorescence images were acquired using a Hamamatsu digital camera. Brightfield images of H&E-stained sections were acquired with an Axiocam 105 color camera. Images were processed and analysed using Zen (v3.5) Blue software (Carl Zeiss MicroImaging GmbH, Germany).

### LPS extraction and silver staining

Overnight bacterial cultures were grown in LB supplemented with appropriate antibiotics and 0.2% glucose. The following day, cultures were diluted 1:100 into fresh LB containing the same antibiotics and 0.2% L-arabinose to induce gene expression from the pBAD promoter. Cultures were incubated at 37 °C with shaking for 3 h. Cultures were then normalised to an OD₆₀₀ of 1.2 in 1 mL volumes and harvested by centrifugation. Supernatants were discarded, and cell pellets were resuspended in 200 μL of 1× Laemmli sample buffer (Bio-Rad), vortexed, and boiled for 5 min. Following heat treatment, 5 μL of proteinase K (20 mg/mL) was added to each sample. Samples were then incubated at 60 °C for 2 h. After incubation, 10 μL of 2-mercaptoethanol was added, and samples were vortexed and boiled again for 5 min. Lysates were centrifuged, and the resulting supernatants were loaded onto a 12 % SDS-PAGE gel for electrophoresis under standard conditions.

Silver staining was performed using the SilverQuest™ Silver Staining Kit (Invitrogen) with an adapted protocol. Gels were rinsed in Milli-Q (MQ) water for 5 min immediately after electrophoresis. Gels were then fixed overnight in 40% isopropanol and 5% acetic acid in MQ water. Fixed gels were washed twice with MQ water (5 min each) and oxidized for 5 min in freshly prepared oxidizing solution containing 0.7% periodic acid, 40% isopropanol, and 5% acetic acid in MQ water. Following oxidation, gels were washed twice in MQ water (15 min each) and then washed in 30% ethanol for 10 min. Gels were sensitised in a solution composed of 30 mL ethanol, 10 mL kit-provided Sensitiser, and 60 mL MQ water for 10 min. After sensitisation, gels were washed once in 30% ethanol (10 min) and once in MQ water (10 min). Staining was carried out by incubating the gels in 100 mL of Staining solution (1 mL kit-provided Stainer in 99 mL MQ water) for 15 min. Gels were developed in 100 mL of Developing solution (10 mL kit-provided Developer, 1 drop of Enhancer, and 90 mL MQ water), until LPS bands became visible. Development was stopped by directly adding 10 mL of kit-provided Stopper into the staining container and gently agitating the gel for 10 min. Gels were then washed in MQ water for 10 min.

Images were acquired using a Chemidoc imaging system (Bio-Rad) and were analysed using ImageLab (v6.1) (Bio-Rad).

### Data analysis and visualisation

No statistical methods were used to predetermine sample size. For ELISAs, two technical replicates were used to calculate experimental means. Statistical significance between two normally distributed groups was performed by Welsh’s *t* test; when two groups were not normally distributed but were lognormally distributed, a lognormal Welch’s *t* test was used. When data were not normally or lognormally distributed (based on D’Agostino-Pearson or Shapiro-Wilk normality tests), a nonparametric Mann-Whitney test was applied. For comparisons involving more than two normally distributed groups, one-way analysis of variance (ANOVA) was used. When assessing more than two normally distributed groups across different timepoints, a two-way ANOVA (for independent measures) or a mixed-effects model (for repeated measures) were performed. When data were not normally distributed (based on D’Agostino-Pearson or Shapiro-Wilk normality tests), a logarithmic transformation was applied and were then analysed by parametric statistical methods (ANOVA). If normality was not achieved by transformation, a nonparametric test was applied (Kruskal-Wallis test followed by Dunn’s multiple comparisons test). Multiple comparisons were corrected using either Dunnett’s or Tukey’s post hoc test, as specified in the figure legends. Statistical analyses were only performed on datasets with a minimum of three or more biological replicates per group. Data visualisation and statistical analyses were performed using Prism (v10.4.1) (GraphPad Software, LLC). A *p* value < 0.05 was considered statistically significant. Statistical tests used for each experiment are detailed in the corresponding figure legends.

## Author contributions

**Audrey Peters:** Conceptualisation, Data curation, Formal analysis, Investigation, Methodology, Validation, Visualisation, Writing – original draft, Writing – review & editing. **Hiba Shareefdeen:** Conceptualisation, Data curation, Investigation, Methodology, Writing – review & editing. **Julia Sanchez-Garrido:** Conceptualisation, Investigation, Methodology, Supervision, Writing – review & editing. **Eli J. Cohen:** Formal analysis, Methodology, Writing – review & editing. **Rémi Denise:** Formal analysis, Methodology, Writing – review & editing. **Joshua L C Wong:** Methodology, Writing – review & editing. **Morgan Beeby:** Supervision, Funding acquisition, Writing – review & editing. **Colin Hill:** Conceptualisation, Funding acquisition, Supervision, Writing – review & editing. **Gad Frankel:** Conceptualisation, Funding acquisition, Supervision, Writing – review & editing.

## Data availability

The ColRes, Eifel1 and Eifel2 bacteriophage genome sequences are available here: https://doi.org/10.5281/zenodo.15724312

All the other data are contained within the main and supplementary material.

## Acknowledgments

We thank Connor Preston for performing histological staining. The Galaxy server used for some calculations is partly funded by the German Federal Ministry of Education and Research BMBF grant 031 A538A de.NBI-RBC and the Ministry of Science, Research and the Arts Baden-Württemberg (MWK) within the framework of LIBIS/de.NBI Freiburg.

